# Profiling skeletal muscle-derived secretome with differentiation and acute contractile activity

**DOI:** 10.1101/2022.02.20.481208

**Authors:** Benjamin Bydak, Taiana M. Pierdoná, Samira Seif, Karim Sidhom, Patience O. Obi, Hagar I. Labouta, Joseph W. Gordon, Ayesha Saleem

## Abstract

Extracellular vesicles (EVs) released from all cells, are essential to cellular communication, and contain biomolecular cargo that can affect recipient cell function. Studies on the effects of contractile activity (exercise) on EVs usually rely on plasma/serum-based assessments, which contain EVs from many different cells. To specifically characterize skeletal muscle-derived vesicles and the effect of acute contractile activity, we used an *in vitro* model where C2C12 mouse myoblasts were differentiated to form myotubes. EVs were isolated from conditioned media from muscle cells, pre-differentiation (myoblasts) and post-differentiation (myotubes), as well as from acutely stimulated myotubes (1hr @ 14V, C-Pace EM, IonOptix) using total exosome isolation reagent (TEI, ThermoFisher, referred to as extracellular particles [EPs]) and differential ultracentrifugation (dUC; EVs). Myotube-EPs (~98 nm) were 41% smaller than myoblast-EPs (~167 nm, p<0.001, N=8-10). Two-way ANOVA showed a significant main effect for size distribution of myotube *vs*. myoblast-EPs (p<0.01, N=10-13). Myoblast-EPs displayed a bimodal size distribution profile with peaks at <200 nm and 400-600 nm, compared to myotube-EPs that were largely 50-300 nm in size. Total protein yield from myotube-EPs was nearly 15-fold higher than myoblast-EPs, (p<0.001 N=6-9). Similar biophysical characteristics were observed when EVs were isolated using dUC: myotube-EVs (~195 nm) remained 41% smaller in average size than myoblast-EVs (~330 nm, p=0.07, N=4-6) and had comparable size distribution profiles as EPs isolated via TEI. Myotube-EVs also had 4.7-fold higher protein yield *vs*. myoblast EVs (p<0.05, N=4-6). Myotube-EPs had significantly decreased expression of exosomal marker proteins TSG101, CD63, ALIX and CD81 compared to myoblast-EPs (p<0.05, N=7-12). Conversely, microvesicle marker ARF6, and lipoprotein marker APO-A1was only found in the myotube-EPs (p<0.05, N=4-12). There was no effect of acute stimulation on myotube-EP biophysical characteristics (N=7), nor on expression of TSG101, ARF6 or CD81 (N=5-6). Myoblasts treated with control or acute stimulation-derived EPs (13 μg/well) for 48hrs and 72hrs showed no changes in mitochondrial mass (MitoTracker Red), cell viability or cell count (N=3-4). Myoblasts treated with EP-depleted media (72hrs) had ~90% lower cell counts (p<0.01, N=3). Our data show that EVs differ in size, distribution, protein yield and expression of subtype markers pre- *vs*. post-skeletal muscle differentiation. There was no effect of acute stimulation on biophysical profile or protein markers in EPs. Acute stimulation-derived EPs did not alter mitochondrial mass nor cell count/viability. Further investigation into the effects of chronic contractile activity on the biophysical characteristics and cargo of skeletal muscle-specific EVs are warranted.

## Introduction

Extracellular vesicles (EVs) were initially discovered in two back-to-back 1983 papers on the recycling of the transferrin receptor in small vesicles released from rat and sheep reticulocytes^1,2^. The term “exosomes” coined by Dr. Rose Johnstone a few years later, refers to what is now known as the smallest family member of EVs^3^. An evolutionary conserved mode of communication, EVs are secreted by all types of cells^4^ and are found in all biological fluids^5^ such as blood^6,7^, saliva^8^, urine^9^, breast milk^10^, human semen^11^ and cerebrospinal fluids^12^. EVs enclose distinct biological cargo that can be modified depending on alterations in the cellular milieu^13^, and can in turn modulate recipient cell function^14^. Classically, EVs are broadly divided into three separate groups depending on their size, and biogenesis pathway^15–18^: exosomes (30-150 nm), microvesicles (100-1000 nm) and apoptotic bodies (1000-5000 nm) and can be isolated using various techniques as we have previously reviewed^19^. EVs mediate the crosstalk between organs and tissues to facilitate coordination and propagation of physiological changes^16^.

In 2006-2007, miRNA was identified within EVs^20,21^, and research interest in EVs skyrocketed once it was established that EVs could transfer nucleic acids between cells. It was not until the late 2000s that studies began to find that EVs could potentiate intercellular communication via transportation of cargo including RNA, protein, membrane receptors and more^17,22–25^. Since then, EVs have been extensively studied as biomarkers of chronic and acute diseases^26–34^, as therapeutic vectors for drug delivery^35–37^, and more recently in physiological contexts.

Skeletal muscle accounts for ~40% of body mass, and serves as an endocrine organ able to secrete a multitude of proteins, lipids, metabolites (i.e. myokines) that are released from muscle upon physical activity, and are essential in mediating some of the systemic effects of exercise^22^. While a plethora of studies have illustrated the endocrine function of skeletal muscle *in vivo*^38^, the first *in vitro* report to show unequivocal evidence for contraction-induced myokine secretion from skeletal muscle cells in culture was only recently published^39^. Myokines can be released through the classical signalling pathway, and also secreted packaged within EVs. In fact, nearly 400 proteins have been identified in EVs from skeletal muscle alone, and many well established myokines have been found inside EVs^16,40–43^. Several papers have demonstrated that skeletal muscle cells are capable of releasing EVs in culture, both from undifferentiated myoblasts and differentiated myotubes^40,41,44^, and that myotube-derived EVs can modulate recipient cell function^42,44^. Given the central role of skeletal muscle in exercise-induced adaptations and whole-body regulation of metabolism, determining the role of exercise-evoked skeletal muscle-derived EVs (Skm-EVs) is critical. While many studies have shown an increase in systemic EVs with exercise in particular^45–49^, identifying the contribution of Skm-EVs is challenging because of several reasons: 1) markers used for skeletal muscle eg. alpha-sarcoglycan^50^, while abundant in muscle, cannot be guaranteed to be expressed in all Skm-EVs, 2) the difficulty in confirming that the Skm-EVs originated from the muscle undergoing contractile activity, and 3) intramuscular injections of fluorescently labelled EVs or genetic manipulation, while an excellent option in rodent students, is not a viable approach for human exercise studies. This underscores the importance of using *in vitro* models of Skm-EVs where contractile activity can be used to mimic exercise, in order to comprehensively characterize Skm-EVs and determine their role in juxtracrine, autocrine and endocrine signalling.

Here, we compared EV characterization as a function of skeletal muscle differentiation and with acute contractile activity. Murine (C2C12) skeletal muscle myoblasts were differentiated into myotubes, and then electrically stimulated using IonOptix C-PACE EM. Conditioned media was collected pre- and post-differentiation to isolate vesicles using total exosome isolation reagent (TEI, ThermoFisher) and differential ultracentrifugation (dUC). We refer to particles isolated via TEI as extracellular particles (EPs), and those via dUC as EVs, as a nod to the lack of specificity of the former when compared to the latter as recommended in the MISEV guidelines^65^. EVs and EPs were isolated and characterized for biophysical properties and expression of marker proteins in accordance with MISEV guidelines^65^. Next, we electrically paced myotubes and isolated EPs from the conditioned media. Electrical stimulation of cultured myotubes is an established method of evoking contractile activity *in vitro*^51^. It has been shown to mimic exercise in C2C12 myotubes^52–54^, and has been used as a surrogate for both acute and chronic exercise models. Lastly, we co-cultured EPs isolated post-acute stimulation with myoblasts and measured changes in mitochondrial content and cell viability.

## Materials and methods

### C2C12 myoblast proliferation and differentiation

C2C12 myoblasts were seeded at 90,000 cells/well in a 6-well dish and grown in fresh DMEM supplemented with 10% fetal bovine serum (FBS) and 1% penicillin/streptomycin (P/S), as previously described^53,55^. After 24 hrs, conditioned media from myoblasts was collected and used for vesicle isolation. Upon reaching 95% confluency, myoblasts were placed in differentiation media (DMEM, 5% horse serum, 1% P/S) for 5 days to get fully differentiated myotubes. Conditioned media from myotubes was collected on day 6 and used for vesicle isolation. Myotubes were electrically stimulated on day 7 as described below to induce acute contractile activity.

### Skeletal muscle EP isolation

Conditioned media was immediately centrifuged at 2,000*x* g to remove cell debris. The pellet was discarded, and supernate used for EP isolation using TEI kit (ThermoFisher, cat #4478359) according to the manufacturer’s instructions and as described before^44^. Modifications to the protocol included nutating conditioned media with 0.5 volume of TEI solution overnight at 4 °C (16 hrs). After nutation, samples were centrifuged at 10,000*x* g for 1 hr (4 °C), washed with 4 ml PBS, and centrifuged again at 10,000*x* g for 1 hr at 4 °C. Pellets were resuspended in 70 μl PBS. The supernate was used for EP-depleted media treatment. Protein concentration of isolated EPs was determined using commercially available BCA protein assay kit (Pierce™, ThermoFisher) as previously described^56^. Protein yield was calculated by multiplying the concentration with total volume of EP isolates.

### Skeletal muscle EV isolation

We used dUC to isolate EVs following the protocol by Théry et al^76^. Media was spun at 300*xg*, 10 min at 4°C to pellet dead cells, followed by *2000xg*, 10 min at 4°C and then 10,000*xg*, 30 min, at 4 °C spins to remove cellular debris and large vesicles, respectively. The supernate was then centrifuged at 100,000*xg* for 70 min at 4°C (Sorvall^™^ MTX 150 Micro-Ultracentrifuge, S58-A fixed angle rotor) to obtain the exosome/small EV pellet. EV pellet was resuspended in 1 mL PBS and centrifuged again at 100,000*xg* for 70 min at 4 °C. The final pellet was resuspended in 50 μL PBS and used for subsequent analysis. Protein concentration of isolated EVs was determined using commercially available BCA protein assay kit (Pierce™, ThermoFisher) as previously described^56^. Protein yield was calculated by multiplying the concentration with total volume of EV isolates.

### Size and zeta analysis

The hydrodynamic diameter (size) and zeta potential of EPs and EVs was characterized by phase analysis light scattering (NanoBrook ZetaPALS, Brookhaven Instruments, Holtsville, NY, USA) instrument in collaboration with Dr. Hagar Labouta’s lab (College of Pharmacy, University of Manitoba). Isolated vesicles were stored for up to 24-48 hrs at 4 °C before characterization. EPs/EVs were diluted 1:75 in PBS and kept on ice until analysis. Each sample underwent 5 runs, each run ~15 seconds, with a dust cut-off set to 40. Size was measured as an intensity averaged multimodal distribution using a scattering angle of 90°, and size bins were used to represent total size intensity within a given size range. Zeta potential analysis was performed using a Solvent-Resistant Electrode (NanoBrook) and BI-SCGO 4.5 ml cuvettes (NanoBrook). For zeta potential, each sample was loaded into the cuvette, and the electrode inserted for phase analysis light scattering to carry out mobility measurements. Values were averaged (irrespective of negative/positive charge) to calculate zeta potential using the Smoluchowski formula from mobility measurements^57^. All measurements were performed in PBS (pH 7.4) at 25 °C.

### Western blotting

For immunoblotting, 20 μg or 50 μg of vesicle protein lysate was denatured with 5% solution of β-mercaptoethanol, incubated at 95 °C for 5 min, then loaded on Mini-PROTEAN^®^ TGX™ Precast Gels (Bio-Rad) for 15 min at 300 V. Proteins were transferred to polyvinylidene difluoride membrane using a Trans-Blot^®^ Turbo™ (Bio-Rad). Once transferred, membranes were washed with TBS-Tween20 (TBST) for 10 min and blocked with 5% skim milk in TBST solution for 2 hrs at room temperature. Membranes was incubated with the primary antibodies against target proteins: rabbit polyclonal anti-TSG 101 (T5701, Sigma-Aldrich Co., 1:200), rabbit polyclonal anti-CD63 (SAB4301607, Sigma-Aldrich Co., 1:1,000), rabbit monoclonal anti-Alix (MCA2493, BioRad, 1:500), mouse monoclonal anti-CD81 (sc-166029, Santa Cruz, 1:200), mouse monoclonal anti-ARF6 (sc-7971, Santa Cruz, 1:200), mouse monoclonal anti-Apolipoprotein A1 (0650-0050, BioRad, 1:200), rabbit polyclonal anti-Cytochrome C (AHP2302, BioRad, 1:200), and mouse monoclonal anti-β-actin (A5441-.2ML, Sigma-Aldrich, 1:5000) in 1% skim milk overnight at 4 °C. Membranes were washed 3x with TBST and incubated with anti-mouse or anti-rabbit IgG horseradish peroxidase secondary antibody (A16017 or A16035, Thermofisher, 1:1,000-10,000) for 1 hr. Membranes were visualized by enhanced chemiluminescence detection reagent (Bio-Rad) and imaged using a ChemiDoc System (BioRad). Some membranes were stripped and re-probed for analysis of target proteins. To do this, membranes were washed 3x with TBST, place in petri dish with Restore^™^ stripping buffer (Thermo Scientific) for 30 min at room temperature, washed with TBST to remove stripping buffer and incubated with primary and secondary antibodies against proteins targets as described above. Band densities of all measured proteins were normalized to the respective Coomassie staining of gels as a loading control.

### Acute stimulation (STIM) of C2C12 myotubes

Electrical stimulation of myotubes has been used to evoke contractile activity and mimic exercise *in vitro* as shown previously^52,53,55^. Prior to STIM, IonOptix C-Dish (stim electrode plates) were sterilized in UV light for 30 min. After sterilization, two electrode plates were loaded into two 6-well dishes with fully differentiated myotubes ready for contractile activity. Once loaded, a single plate (designated as STIM) was connected to the stim machine (IonOptix C-Pace EM), whereas the control plate was not connected to the IonOptix C-Pace EM. Stimulation was performed at 14 V, 1 Hz, for 1 hr, while both plates were incubated at 37 °C. Immediately after STIM, media was collected, and EP isolation performed as described above using TEI.

### Treatment of C2C12 myoblasts with EPs isolated from control and STIM myotubes

90,000 myoblasts/well were seeded in 6-well plates in 2 ml/well of growth media. 6.67 μg/ml EPs (total 13 μg EPs/well) or 1 ml EP-depleted media from control or STIM myotubes was added to myoblasts when cells were at 80-90% confluency. Cells were treated for 48 hrs and 72 hrs at 37 °C. After treatment, media was discarded and treated myoblasts collected for MitoTracker Red CMXRos staining, cell count or viability assays as described below.

### MitoTracker staining

MitoTracker Red CMXRos (Cell Signaling, #9082) was first prepared to a concentration of 0.1 M in 1X PBS. 60 μl of 0.1 M of MitoTracker Red CMXRos was then mixed in 12 ml of growth media to prepare staining solution. Cells were then washed twice with 1X PBS, stained with 800 μl of staining solution, and incubated at 37 °C for 30 min. Following incubation, cells were washed 2x with PBS and covered with 1 ml of growth media before imaging using epifluorescence microscopy (Zeiss Axiovert 200).

### Cell count and viability assay

Cells were washed twice with 800 μl PBS, trypsinized with 400 μl of trypsin, and incubated for 3 min at 37 °C. 1 ml of growth media was added, and cells were centrifuged at 1000*x* g for 5 min to pellet the cells. Pelleted cells were stained with a 1:1 dilution of 0.4% Trypan blue solution (Sigma, cat #T8154), then counted with a hemocytometer (Hausser Scientific Bright-Line Hemacytometer). Total number of cells were counted and expressed per ml of growth media. Cell viability was obtained by dividing the number of live cells (not stained by the trypan) by the total amount of counted cells.

### Statistical analysis

All data were analyzed using an unpaired Student’s t-test. Size distribution, and cell count was assessed using a two-way ANOVA with Bonferroni post-hoc correction for multiple comparisons. Significance was set at p<0.05. All data are presented as mean ± standard error. Statistical analysis was performed using GraphPad Prism software (version 8.4.2).

## Results

### Isolated vesicles differ in size, protein yield and expression of vesicle subtype markers pre- and post-differentiation

Average myotube-EP size (98 nm) was 41% smaller than myoblast-EPs (167 nm, p<0.001, **Fig. 1A**). The smallest particle size was 63 nm in myotube-EPs *vs*. 102 nm in myoblast-EPs, while the maximum size observed was 130 nm in myotube-EPs *vs*. 206 nm in myoblast-EPs. There was no difference in zeta potential between the two groups (**Fig. 1B**). A two-way ANOVA on size distribution between myoblast-EPs and myotube-EPs showed a significant main effect for cell type (p<0.05, **Fig. 1D**). Myoblast-EPs show a bimodal EV size distribution pattern with two distinct peaks in expression, one at <200 nm, and second at 400-600 nm (**Fig. 1D**). In contrast, myotube-EPs are largely enriched with 50-300 nm sized particles (**Fig. 1D**). Total protein yield from myotube-EPs was ~15-fold higher than myoblast-EPs (p<0.001, **Fig. 1C**). To compare our results with the gold-standard method of EV isolation, we next used dUC to isolate EVs. Average EV size, distribution profile, and zeta potential were not statistically different between myotube-EVs *vs*. myoblast-EVs (**Fig. 2A, B, D**). Despite this lack of statistical significance, we observed the same trends. Myotube-EVs (~195 nm) remained 41% smaller in average size than myoblast-EVs (~330 nm, p=0.07, N=4-6, **Fig. 2A**). The minimum EV size was 138 nm for myotube-EVs, and 184 for myoblast-EVs. The maximum EV size was 290 nm for myotube-EVs *vs*. 570 nm for myoblast-EVs. Further, myoblast-EVs displayed a biomodal size distribution with enrichment of <200 nm, and 400-600 nm sized vesicles (**Fig. 2D)**. Myotube-EVs conversely were enriched with 150-300 nm sized EVs (**Fig. 2D**). Myotube-EV protein yield was 4.79-fold higher than myoblast-EV protein yield (p<0.05, **Fig. 2C**), albeit overall protein yield was 20-40x lower in EVs isolated via dUC (**Fig. 2C**), when compared to the TEI method of EP extraction (**Fig. 1C**).

**Figure 1.**
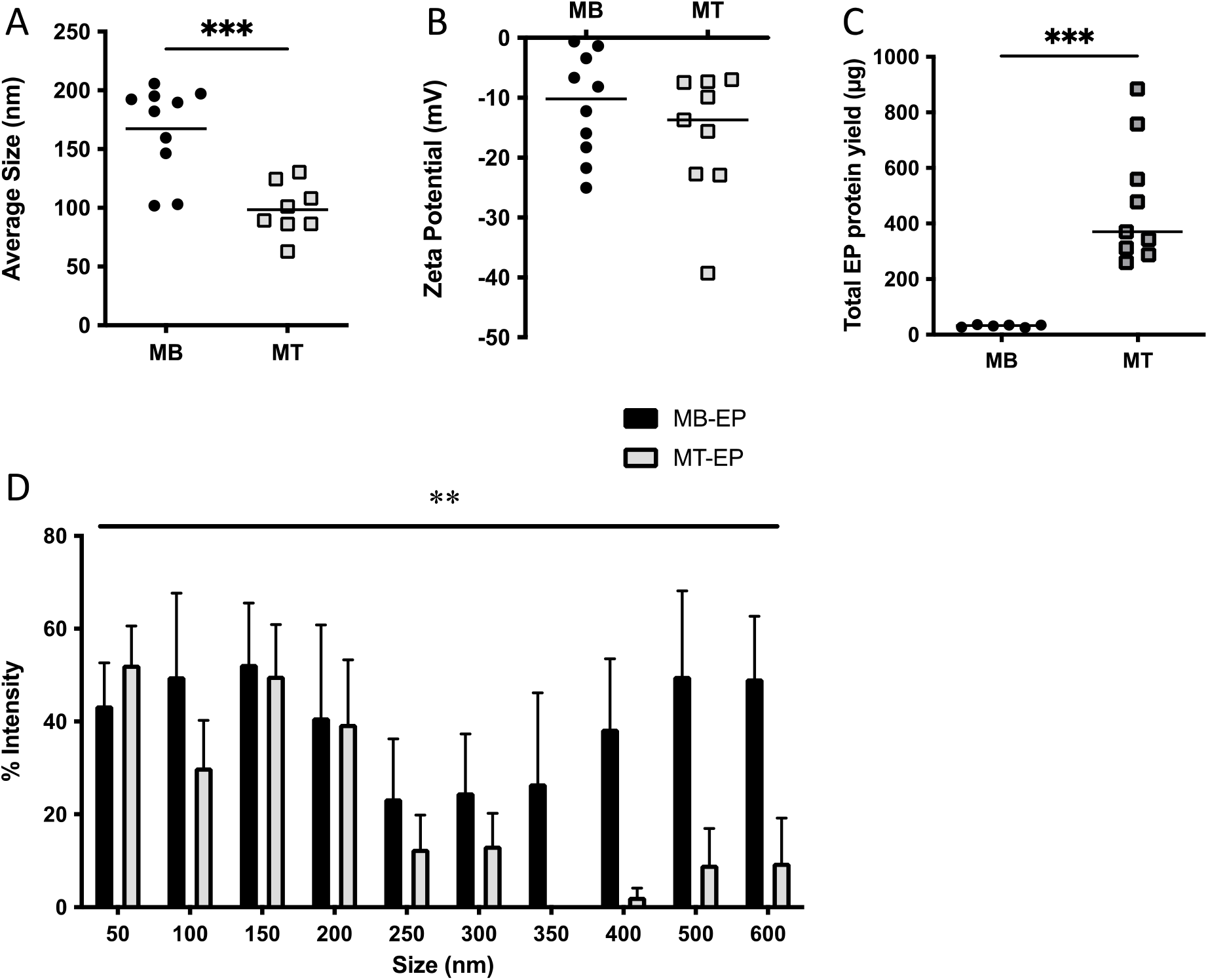
Changes in EP biophysical characteristics with skeletal muscle differentiation. (A) C2C12 myotube (MT) EPs were 41% smaller than myoblast (MB) EPs (***p<0.001, N=8-10). (B) Zeta potential remained unchanged between EPs from myoblasts *vs*. myotubes, N=9-10. (C) Total protein yield from myotube-EPs was ~15-fold higher than myoblast-EPs (***p<0.001 N=6-9). (D) Two-way ANOVA showed a significant main effect for myotube-EPs *vs*. myoblast-EPs (**p<0.01, N=10-13). Myoblast-EPs display a bimodal size distribution profile with a peak for EPs <200 nm and again at 400-600 nm, compared to myotube-EPs that were enriched with 50-300 nm sized particles. EPs were isolated using Total Exosome Isolation Reagent (ThermoFisher, cat# 4478359). All data were analyzed using an unpaired Student’s t-test, except in Fig. 1D where a 2-way ANOVA was used. Data are expressed as scatter plots showing mean, or bars with mean ±standard error.

**Figure 2.**
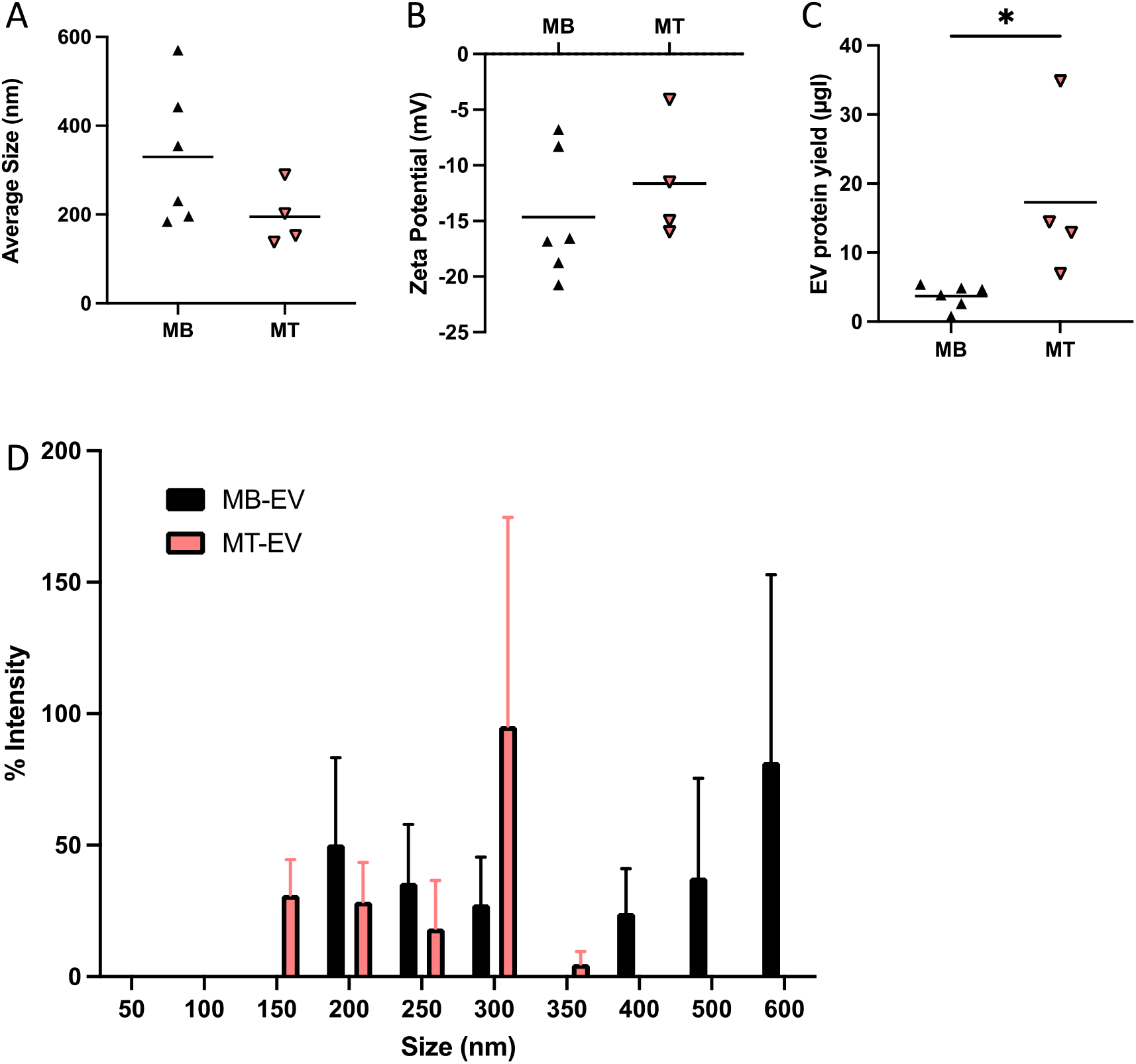
Differences between myoblast (MB) vs. myotube (MT) EV biophysical characteristics when isolated using differential ultracentrifugation (dUC). C2C12 myoblast- and myotube-EVs isolated via dUC showed no difference in (A) average EV size or (B) zeta potential (N=4-6). (C) Myotube EV protein yield was 4.79-fold higher than myoblast EV protein yield (*p<0.05, N=4-6). (D) Myoblast EVs display a bimodal size distribution profile with a peak at 200 nm and again at 600 nm, compared to myotubes EVs that were largely enriched with EVs between 150-300 nm (N=4-6). All data were analyzed using an unpaired Student’s t-test, except in Fig. 2D where a 2-way ANOVA was used. Data are expressed as scatter plots showing mean, or as bar graphs with mean ± standard error.

Next, we compared the expression of protein markers commonly associated with small EVs or exosomes i.e. TSG101, Alix, CD81 and CD63, and medium/large EV (i.e. microvesicle) marker ARF6 to characterize EPs in compliance with MISEV guidelines^65^. Myotube-EPs had significantly decreased expression of small EV protein markers (TSG101, CD63, ALIX and CD81), often by several orders of magnitude compared to myoblast-EPs (p<0.05, **Fig. 3A** and **3B**). Conversely, APO-A1 (lipoprotein) expression and ARF6 expression were highly enriched in myotube-EPs *vs*. myoblast-EPs (p<0.05, **Fig. 3A** and **3B**). Lastly, cytochrome c (mitochondrial marker used to identify medium/large EVs) and beta-actin (housekeeping control) were barely expressed at quantifiable levels in myotube-EPs compared to myoblast-EPs (p<0.05, **Fig. 3A** and **3B**).

**Figure 3.**
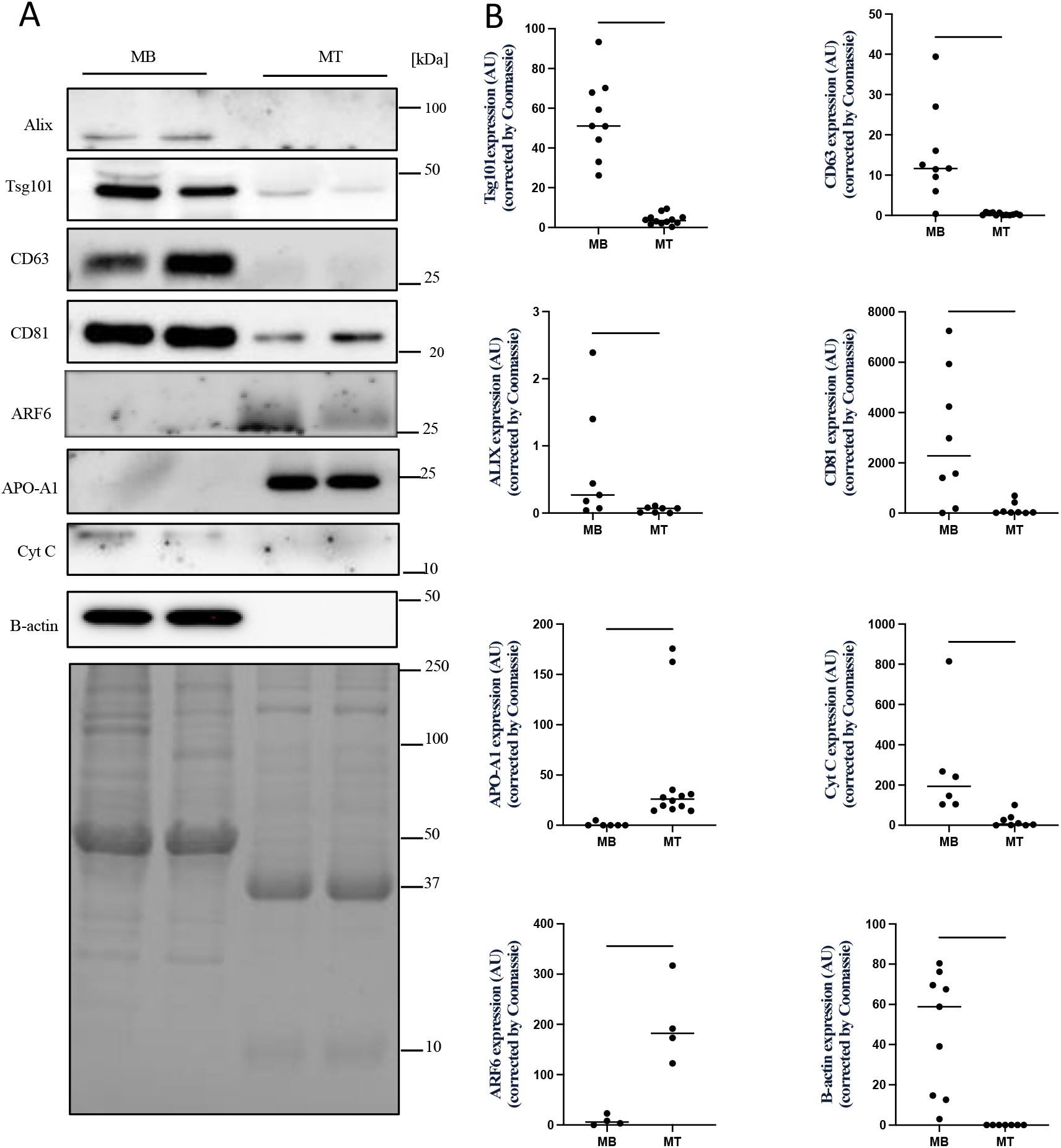
Effect of skeletal muscle cell differentiation on the expression of proteins related to EV subtypes. (A) Representative immunoblots showing equal amounts (20 μg) of myoblast (MB) or myotube (MT) EP protein lysates were subjected to 12% SDS-PAGE. Coomassie blue gel staining was used as a loading control. (B) Quantification of immunoblot analysis showing the expression of different markers of small and med/large EVs and non-EV co-isolates. Expression of small EV marker proteins TSG101, ALIX, and tetraspanins CD63 and CD81, were enriched by several orders of magnitudes in myoblast-EP preparations compared with myotube-EPs (*p<0.05, **p<0.01, ***p<0.001, ****p<0.0001, N=7-12). Conversely, levels of ARF6, a microvesicle marker, and APO-A1, a lipoprotein non-EV co-isolate, were only expressed in myotube-EPs when compared with myoblast-EPs (*p<0.05, **p<0.01, N=4-12). Cytochrome c and beta-actin were barely expressed in myotube-EPs compared to myoblast-EPs (**p<0.01, N=6-9). Data were analyzed using an unpaired Student’s t-test, are expressed as scatter plots showing the mean.

### Acute stimulation does not affect vesicle size, zeta potential, protein yield or expression of vesicle subtype protein markers

To quantify the effects of acute contractile activity on myotube-EPs, fully differentiated myotubes were electrically paced at 14 V (1 Hz) for 1 hr. Control myotubes were plated with the stimulatory electrode plate, but did not receive stimulation. Immediately after stimulation, we collected conditioned media and performed EP isolation as described earlier using TEI. We found no difference in average size (**Fig. 4A**), zeta potential (**Fig. 4B**), total protein yield (**Fig. 4C**) or size distribution in control *vs*. stimulated conditions (**Fig. 4D**). We evaluated the expression of small EV markers with stimulation. Due to the extremely low levels of TSG101 observed in myotube-EPs (**Fig. 3**), we decided to run Western blots with higher total protein content per lane (50 μg) for these experiments (**Fig. 5**). We found no difference in the expression of ARF6, TSG101 or CD81 in control *vs*. stimulated conditions (**Fig. 5A** and **5B**).

**Figure 4.**
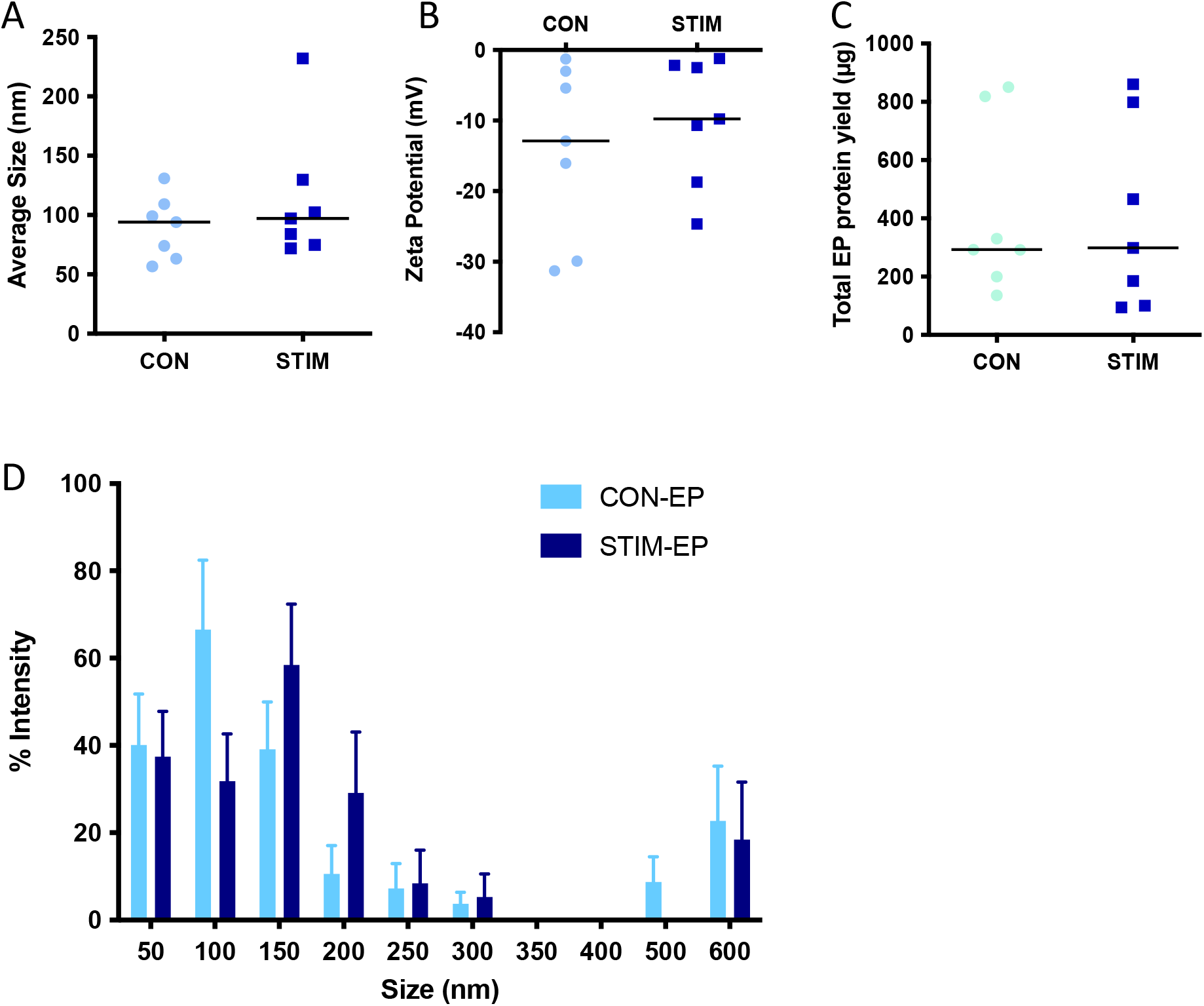
Effect of acute stimulation on EP size, zeta potential, and protein yield. C2C12 myotubes were electrically contracted using an IonOptix EM-PACE for 1 hr at 14 volts and EPs extracted from conditioned media. (A) Average size, (B) zeta potential and (C) protein yield remained unchanged between control (CON) and acutely stimulated (STIM) myotube-EPs (N=7). (D) EP size distribution was also relatively similar between both CON and STIM (N=7). EPs were isolated using TEI. Data were analyzed using an unpaired Student’s t-test except for panel D which was analyzed by 2-way ANOVA. Data are expressed as scatter plots with means or bar graphs with mean ± standard error.

**Figure 5.**
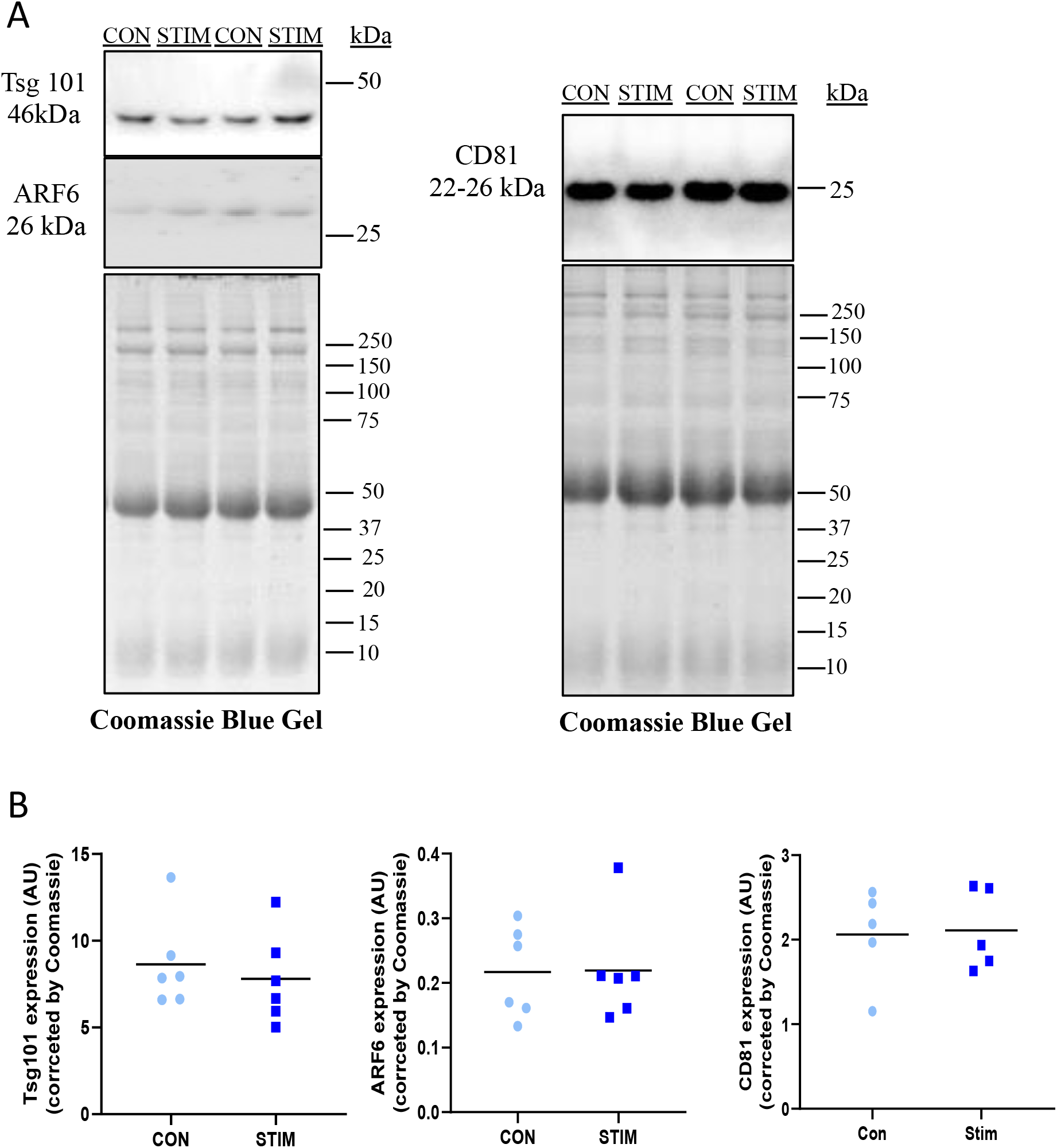
Effect of acute stimulation on expression of EV subtype protein markers. Equal amounts (50 μg) of control (CON) *vs*. acutely stimulated (STIM) myotube-EP lysates were subjected to 12% SDS-PAGE. Coomassie blue-stained gels were used as a loading control. (A) Representative immunoblots and (B) quantification of data shows no difference in the expression of exosomal markers CD81 and TSG101, nor in the content of microvesicle marker ARF6 in EPs lysates CON *vs*. STIM myotubes (N=5-6). Data were analyzed using an unpaired Student’s t-test, are expressed scatter plots with mean.

### Effect of vesicles collected post-stimulation on mitochondrial content, cell count and cell viability

We performed EP co-culture experiments to measure the potential of EPs after an acute stimulation to deliver an adaptive metabolic response in other cells. We seeded C2C12 myoblasts at 90,000 cells/well in a standard 6-well plate. Each well was incubated with 13 μg (6.67 μg/ml) of freshly isolated EPs from control or stimulated myotubes for 48 hrs and 72 hrs. This dosage was empirically determined as shown previously^42,58,59^. After treatment, we stained the cells for mitochondrial content with 0.1 mM MitoTracker CMXRos for 30 min. Representative images at 10X are shown for myoblasts treated with control and stimulated myotube-EPs for 48 hrs and 72 hrs (**Fig. 6A**). No significant difference was found between the mitochondrial staining corrected by total nuclei count at either 48 hrs or 72 hrs time point (**Fig. 6B**). To determine effect on cell viability, control or stimulated EP-treated myoblasts were counted using a trypan blue exclusion assay after 72 hrs treatment. Cell viability was not affected with control or stimulated myotube-EP treatment (**Fig. 6C**). There was also no significant difference in total cell count at 72 hrs with stimulated myotube-EP *vs*. control treatment (**Fig. 7**). However, myoblasts treated for 72 hrs with conditioned media that was depleted of EPs (EP-dep, 1 ml) displayed a ~90% decrease in cell count (**Fig. 7**), irrespective of whether they were treated with control or stimulated EP-dep media.

**Figure 6.**
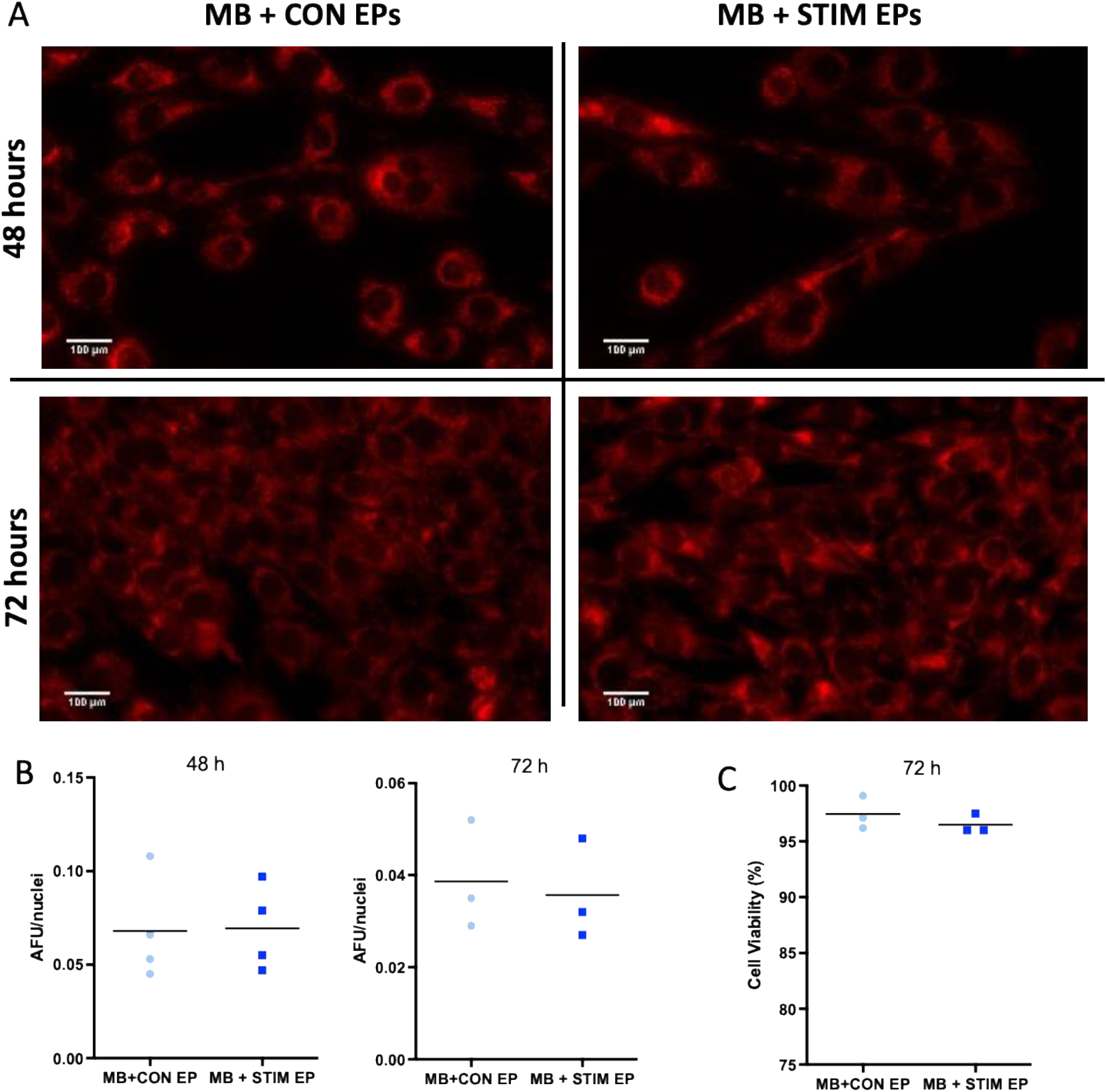
Mitochondrial content and cell viability in myoblasts (MB) treated with EPs from CON *vs*. STIM myotubes. Myoblasts were treated with EPs (13 μg/well) isolated from conditioned media from control (CON) *vs*. acutely stimulated (STIM) myotubes for 48 hrs and 72 hrs and changes in mitochondrial mass measured. (A) Representative fluorescent images of myoblasts stained with MitoTracker Red CMXRos after 48 hrs and 72 hrs of EP treatment from CON and STIM myotubes taken at 10X mag, Scale bar =100 μm. (B) Quantification of fluorescent images showed no change in mitochondrial content with STIM myotube-EP treatment. Images were normalized to nuclei count by dividing total fluorescence of each image with number of nuclei and expressed as arbitrary fluorescent units (AFU) per nuclei (N=3-4). (C) Cell viability, determined by trypan blue exclusion and expressed as % viable cells remained unchanged between CON *vs*. STIM treated myoblasts (N=3). Data were analyzed using an unpaired Student’s t-test, and are expressed as scatter plots with mean.

**Figure 7.**
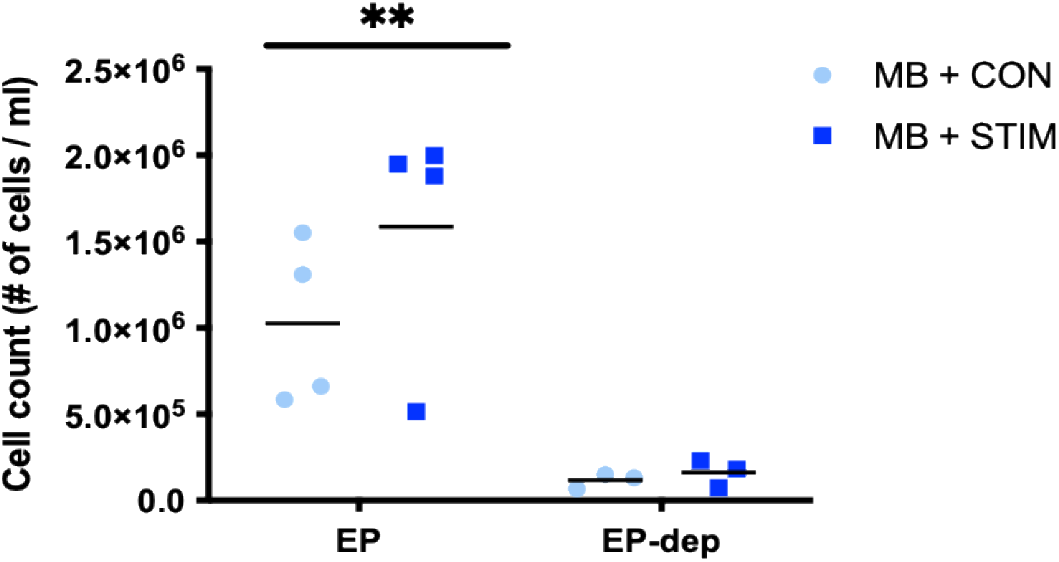
Cell count in myoblasts (MB) treated with EPs or EP-depleted media from CON *vs*. STIM myotubes. 90,000 cells/well were seeded in 6-well plates and left to adhere for four hours. Cells were treated with 13 μg/well EPs or with EP-depleted media (EP-dep, 1ml) conditions from control (CON) *vs*. acutely stimulated (STIM) myotubes for 72 hrs. Total number of cells were counted using a haemocytometer and expressed per ml of media. Cell count was ~90% lower in myoblasts treated with EP-dep conditions from either CON or STIM myotubes (**p<0.01, main effect of EP *vs*. EP-dep, N=3-4).

### Effect of fetal bovine serum (FBS) and horse serum (HS) on vesicle preparations

C2C12 myoblasts were cultured in growth media (GM) containing 10% FBS as previous research has shown that growing cells in exosome-depleted FBS negatively affects cell growth^60^. Similarly, myotubes were placed in differentiation media (DM) supplemented with 5% HS that was not EV-depleted. Given that both FBS and HS can contain EVs from source that can confound the data, we isolated EPs from GM and DM only, and compared them to the EPs isolated from myoblasts and myotubes, respectively. Media was placed in 6-well plates with no cells, and incubated at 37 °C for 24 hrs to mimic the myoblast/myotube-conditioned media acquisition procedure. EP size, zeta potential and protein yield in particles isolated from GM and DM only conditions is shown in **Table 1**. We compared GM-EP biophysical characteristics to myoblast-EPs, and DM-EP biophysical characteristics to myotube-EPs using unpaired Student’s t-tests. While EP size and zeta potential was not different, there was a 4.7-fold increase in protein yield in GM-EPs *vs*. myoblast-EPs (p<0.05, **Table 1**). DM-EPs were 68% larger in average size, and had 72% less protein yield compared to myotube-EPs (p<0.05, **Table 1**).

**Table 1.**
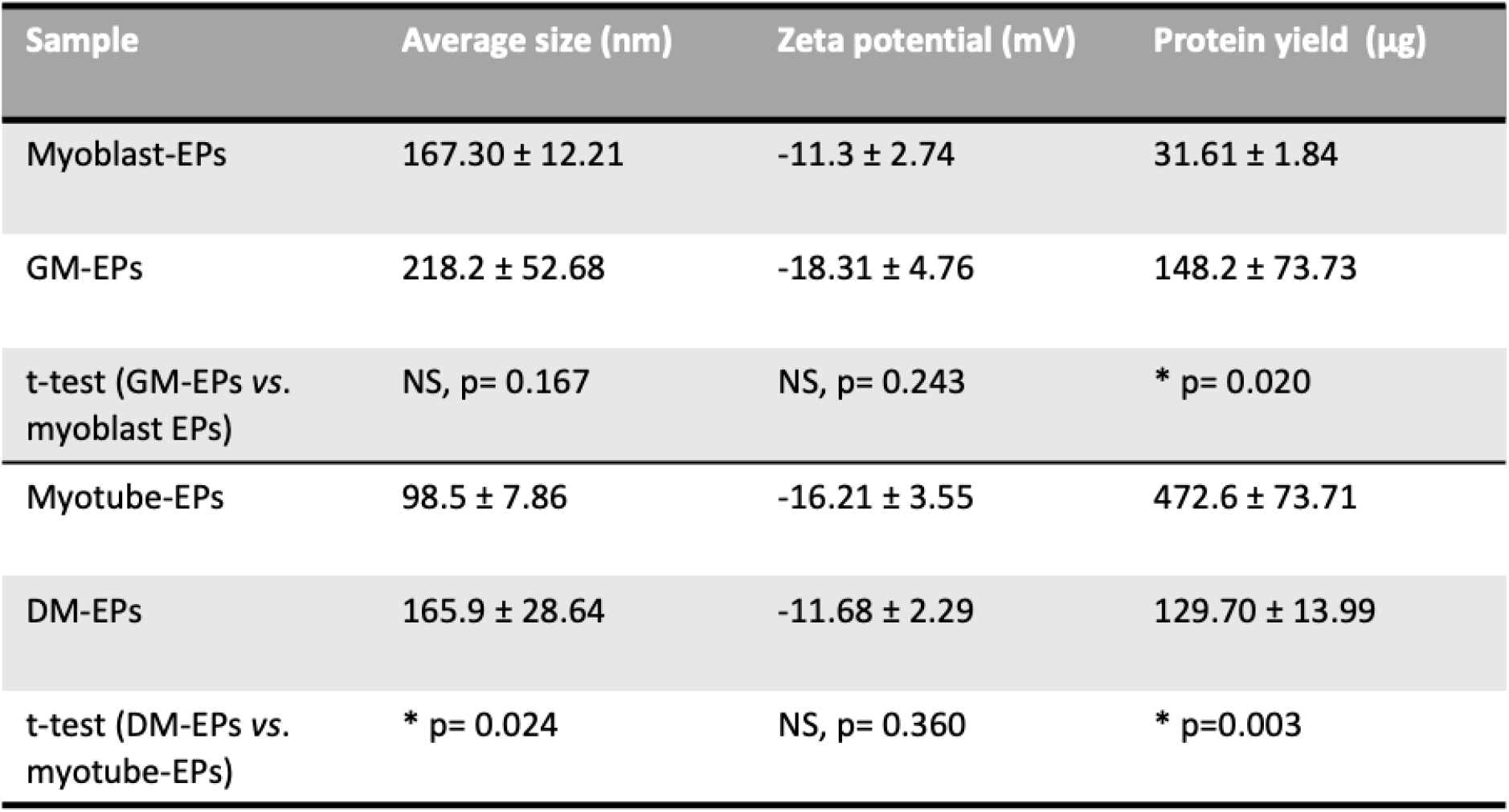
Average size, zeta potential, and protein yield of EPs isolated from myoblasts (MB) and myotubes (MT), and their respective media-only conditions. C2C12 myoblasts were cultured in growth media (GM) made from DMEM containing 10% fetal bovine serum (FBS) and 1% antibiotics. To differentiate myoblasts into myotubes, media was switched to differentiation media (DM): DMEM supplemented with 5% horse serum (HS), 1% antibiotics. Both FBS and HS can contain EVs from bovine and horse serum, respectively. To evaluate the confounding effect of the serum (if any), EPs were isolated using TEI from GM only and compared to myoblast-EPs, and from DM only and compared to myotube-EPs (N=3-6, *p< 0.05). Results were analyzed using an unpaired Student’s t-test. Data are expressed as mean ± standard error.

## Discussion

Our results indicate that skeletal muscle differentiation affects the vesicle size, protein yield and expression of subtype protein markers. Importantly the differences in vesicle biophysical characteristics were observed with both a crude method of particle isolation (TEI), as well as with the gold standard method of EV isolation, dUC^19^. Acute contractile activity had no affect on the biophysical properties of EPs and culturing myoblasts with control *vs*. stimulated myotube-EPs did not affect mitochondrial mass nor cell viability/count.

Previous work describing average size of myoblast and myotube EVs are largely consistent in their description of both cells releasing primarily small EVs^40,41,61,62,77^. Forterre et al. (2014)^41^ reported myoblast- and myotube-EVs had similar average EV size, with no particles over 500 nm being found. However, in this study the authors used a 0.2 μm filter before dUC so that would have removed any particles over 300 nm. Similarly, others using dUC and 0.2 μm filtration have reported average myoblast-EV size ranges between 50-100 nm^61,62^. The EVs in these studies were analyzed by either DLS^41^, and/or nanoparticle tracking analysis (NTA) and transmission electron microscopy (TEM) analysis^61,62^. We observed myoblasts display a bimodal expression of EV sizes (<200 nm, and 400-600 nm), consistently in both TEI-derived EPs and dUC-derived EVs, without filtration in either method, and analyzed biophysical properties using DLS. Thus, it is likely that the discrepancy in sizes can be explained by the method of vesicle isolation and/or characterization. Further, in our study vesicle size was likely smaller in EPs *vs*. EVs, as TEI precipitates exomeres as well as other non-EV co-isolates that can affect the size profile as detailed in MISEV guidelines^65^. While dUC is the preferred and no doubt the orthogonal approach for isolation of EVs compared to TEI-based precipitation, our results show similar trends in size and protein yield between the two methods which will be important for laboratories that do not have access to high speed ultracentrifugation infrastructure.

We found that myotube-EPs expressed lower expression of small-EV markers (Alix, TSG10, CD63, CD81), and higher expression of microvesicle marker (ARF6) and non-EV co-isolate (APO-A1, lipoprotein marker) compared to myoblast-EVs. Romancino et al. (2013)^40^ have postulated that myotube-EVs *vs*. myoblast-EVs may have different origins and that myotube-EVs are likely preferentially released through plasma membrane budding (microvesicles). This supports the increased ARF6 expression in myotube-EPs in our study. Previous proteomic analysis on myotube-EVs has indicated they contain less endosomal/lysosomal CD63, as opposed to CD81 or CD9 tetraspanins that are also expressed at the plasma membrane^41^. In line with this, our study shows myotube-EPs are CD63- and CD81+. However, we did not observe higher Alix or CD81 expression in myotube-EPs compared to myoblasts, as reported by others^40,41^, nor similar levels of TSG101 expression^41^, likely due to differences in methodology as noted above. As noted by others, expression of small EV/microvesicle markers proteins is cell-specific as well as dependent on isolation methodologies. Hence, expression alone cannot be used to precisely confirm vesicle origin or sub-type^63–65^.

Our results showed that myotube-EPs contained 15-fold more protein than myoblast-EPs when obtained using TEI. This difference between pre- and post-differentiated skeletal muscle cells remained in the dUC-derived EVs, albeit the magnitude was markedly reduced: myotube-EVs had 4.79-fold higher protein yield than myoblast-EVs. It is well known that TEI allows for high recovery but low specificity of separated vesicles, whereas dUC permits intermediate to high specificity but low recovery of vesicles^65^. Consequently, we expected absolute protein yield to be higher in TEI-derived EPs, likely as TEI pulls down non-EV co-isolates and proteins. Given that myotube-EPs expressed significantly elevated levels of APO-A1, protein contamination from non-EV sources is likely amplifying protein yield in this method. Another source of contamination can be proteins from serum in the conditioned media preparations. This protein contamination from media can be present in both myoblast-EP and myotube-EP isolates. Although differentiating myotubes in serum-depleted media 24 hrs prior to vesicle isolation has been used to mitigate this problem^66^, growing myoblasts in exosome-depleted serum has been shown to negatively affect their proliferation and differentiation^67^, indicating that an exosome- or serum-depleted approach may alter cell behaviour and may not be advisable given the experimental context. In this study we chose not to use vesicle-depleted serum, and instead established a baseline of EP biophysical properties expected in media only conditions. Encouragingly, we found that EPs isolated from growth or differentiation media only, had significantly different protein yields compared to our myoblast and myotube-EP preparations, respectively. Differentiation media-derived EPs were also larger sized than myotube-EPs. This indicates that the confounding effect of media-derived proteins/particles on skeletal muscle-derive EP preparations in our study was limited. Lastly, the difference in protein yield between myotube-EVs *vs*. myoblast-EVs remained consistent when we used dUC to isolate the vesicles. Increased myotube-EV total protein content has also been reported by others previously^40^. This suggests that a significant increase in protein yield in myotube-EVs compared with myoblasts can be due to higher EV concentration, or enhanced EV protein cargo levels, or a combination of both. We were not able to ascertain EV concentration due to lack of access to NTA/TRPS infrastructure at this point.

To our knowledge, this is the first characterization of EPs from myotubes after acute stimulation. Electrical pulse stimulation has been used extensively as a method of mimicking exercise and evoking the downstream effects associated with acute and chronic contractile activity in myotubes^68^. That average EP size did not change after acute stimulation is not surprising. No change in average vesicle size has been reported after acute aerobic exercise in rats, or humans performing treadmill or cycle ergometer exercise^69–71^. Furthermore, presence of small EVs after acute exercise in humans have been reported to be 50-300 nm in size^46^, in congruence with our results. However, each of the aforementioned studies reported an increase in small EV concentration after acute exercise. While we could not measure EV concentration in either the control or stimulation groups, our results show no significant differences in EV protein yield after acute stimulation. Furthermore, expression of small EV (TSG101, CD81) or microvesicle (ARF6) marker proteins did not change after acute stimulation. Previously, 1 hr stimulation of C2C12 myotubes has been shown to induce release of IL-6, a well-known myokine^39^, as well increased cellular levels of proliferator activated receptor gamma coactivator 1 alpha (PGC-1α) and AMPK^72^, indicating that 1 hr of electrical stimulation is sufficient to induce myokine release in C2C12 myotubes, and evoke the cellular signalling milieu post-contractile activity. It is likely that a higher intensity, or longer stimulation period would elicit changes in EV size or biophysical properties but that needs to be experimentally determined. Additionally, acute exercise has been shown to increase CD61+ and CD81+ EVs^46^ in human participants. Since our results showed no difference in CD81 expression, this may indicate that the source of increased CD81 expression is species-specific, or lies outside of skeletal muscle-derived EVs. Similarly, while acute exercise in known to increase circulating levels of microvesicles, these are predominantly platelet-derived^73^, which supports our observations showing no difference in ARF6 expression between control and stimulated myotube-EPs.

Since EV cargo can be taken up by recipient cells, and can regulate their fate^16^, we decided to evaluate the biological action of acute stimulation-derived EPs. To do so, we co-cultured myoblasts with control *vs*. stimulated myotube-EPs. After 48 hrs and 72 hrs EV treatment, we noted no significant effect of control *vs*. stimulated myotube-EPs on increasing mitochondrial mass as measured by MitoTracker staining. Increased mitochondrial content is a hallmark adaptation of contractile activity/exercise. Within hours of contractile activity, a cascade of reactions occur whereby PGC-1α is upregulated, which results in the downstream increase of nuclear respiratory factor 1 (NRF-1), and mitochondrial transcription factor A (Tfam), leading to a co-ordinated increase in both nuclear and mitochondrial proteins^74^. Within days to weeks of repeated contractile activity, mitochondria can populate the previously exercised muscle cell^75^. Since we measured mitochondrial content by mitochondrial staining, we did not assess if the aforementioned proteins associated with mitochondrial biogenesis could have been induced. This gap can be addressed in future studies. It is also likely that while treatment with stimulated myotube-EPs may trigger the protein signalling cascade upstream of mitochondrial biogenesis, a single dose is not powerful enough to elicit changes in organelle synthesis. EP treatment from chronically stimulated myotubes could further elucidate the potential of skeletal muscle-derived vesicles to transmit and deliver an exercise response in non-stimulated cells. A growing number of studies have shown that skeletal muscle-derived EVs can have important paracrine effects that affect physiological function (e.g. myogenesis), as well as play a crucial rule in chronic diseases like insulin-resistance, as reviewed comprehensively by others^78^. This warrants further investigation into the biogenesis, uptake, and downstream biological activity of skeletal muscle-derived EVs particularly with chronic exercise.

We acknowledge the limitations in our current study. Polyethylene glycol (PEG)-based TEI kit based precipitation of EPs is known to be a high yield, low purity method of vesicle isolation^65^. Proteins from the media may be pulled into the isolated EP pellet and may mask/confound the true difference in EP preparations. We only measured mitochondrial staining as a proxy for mitochondrial mass/content. Evaluating changes in signalling cascades and protein expression/translocation that are upstream of an increase in mitochondrial content could provide deeper insight into whether stimulation derived myotube-EPs can evoke a metabolic response in treated cells. Lastly, experiments with longer treatment time, higher dosage, and duration of contractile activity i.e. EPs derived post-chronic contractile activity, can provide a better understanding of the role of skeletal muscle-derived EVs in conferring any of the adaptive metabolic changes commonly associated with contractile activity.

## Author Contributions

B.B., T.M.P, S.S. K.S. and P.O.O. performed experiments in the current study, analyzed data, and created figures. B.B., K.S. and A.S. helped draft and revise the manuscript. H.I.L. and J.R.G. provided technical and theoretical expertise to complete the work. All authors were involved in manuscript revisions. A.S. designed the project, and helped synthesize data, create figures, write, and edit the manuscript. A.S. is the corresponding author and directly supervised the project. All authors have read, edited and agreed to the published version of the manuscript.

## Acknowledgments

We would like to thank Oluwaseyi Adegbasan and Carlynn Davidson for technical assistance during the study.

